# MoGAAAP: A modular Snakemake workflow for automated genome assembly and annotation with quality assessment

**DOI:** 10.1101/2025.08.26.672321

**Authors:** Dirk-Jan M. van Workum, Kuntal K. Dey, Alexander Kozik, Dean Lavelle, Dick de Ridder, M. Eric Schranz, Richard W. Michelmore, Sandra Smit

## Abstract

With the current speed of sequencing, there is a desire for standardised and automated genome assembly and annotation to produce high-quality genomes as input for comparative (pan)genomics. Therefore, we created a convenience pipeline using existing tools that creates annotated genome assemblies from HiFi (and optionally ultra-long ONT and/or Hi-C) reads for a set of related accessions as well as a related reference genome. Our pipeline is species-agnostic and generates an extensive quality assessment report that can be used for manual filtering and refinement of the assembly and annotation. It includes statistics for individual completeness and contamination assessments as well as a concise pangenome view.

The pipeline is implemented in Snakemake and available with a GPLv3 license at GitHub under github.com/dirkjanvw/MoGAAAP and at Zenodo under doi.org/10.5281/zenodo.14833021.

## Introduction

With the increase in DNA sequencing throughput, read length and base-calling accuracy, the bottleneck in genomics research has shifted from data generation to data analysis. Sequencing the first human genome took many years, whereas a human genome can currently be sequenced in just a few days and assembled in half a day approximately. Pacific Biosciences HiFi and Oxford Nanopore (ONT) long-read sequencing technologies now make it possible to generate whole genome assemblies for many genotypes per species. Building on existing high-quality reference genomes, efficient workflows are needed to create annotated assemblies for these additional genotypes of the same species.

The genome assembly process itself has improved over the years with a focus on novel algorithms for assembly and integration of multiple types of input data. For complex eukaryotic genomes, subsequent structural and functional annotation is still a laborious process; accurate, *de novo* annotation of a genome assembly (also involving repeat masking) typically takes up to a week for computation alone, with additional manual evaluation and filtering. However, to annotate successive genome assemblies for a species, faster approaches, such as the transfer of annotations from reference genomes and *ab initio* predictions based on machine learning, are available.

Here we address the need for both 1) increasing the speed of creating annotated genome assemblies while decreasing the effort required, and 2) standardising the output, necessary for downstream comparative (pan)genomics. To this end, we implemented MoGAAAP, an automated workflow using a modular Snakemake pipeline that fully automates the processes of assembly, provisional annotation and quality assessment (QA) for any diploid eukaryotic organism. MoGAAAP does not aim to create a perfect, publication-quality genome assembly but rather to provide a high-quality foundation for further refinement, if required. Also, due to its modular nature, MoGAAAP can be used for only assembly, annotation or QA of already existing genomes in any combination. The pipeline creates a standardised QA report which provides the user with suggestions for additional (manual) curation and filtering of the input reads, assembly and annotation. Such automation of these processes will significantly increase the speed of exploring genome repertoires of diverse eukaryotic species.

## Implementation

We created a generalised modular Snakemake (≥v8.0) pipeline [Köster and Rahmann, 2012] that takes as input HiFi (and optionally ONT and/or Hi-C) reads as well as a chromosome-level, annotated reference genome, and creates an annotated draft genome assembly, including a QA report (Fig. 1). The overall process takes about two days for a human genome; however, the exact runtime depends on genome size and available computational resources. Because of the modularity of MoGAAAP, the starting point of an analysis can also be an already created pseudomolecule-level assembly (with or without annotation). This means that MoGAAAP can also be used for QA of existing genomes.

**Fig. 1:**
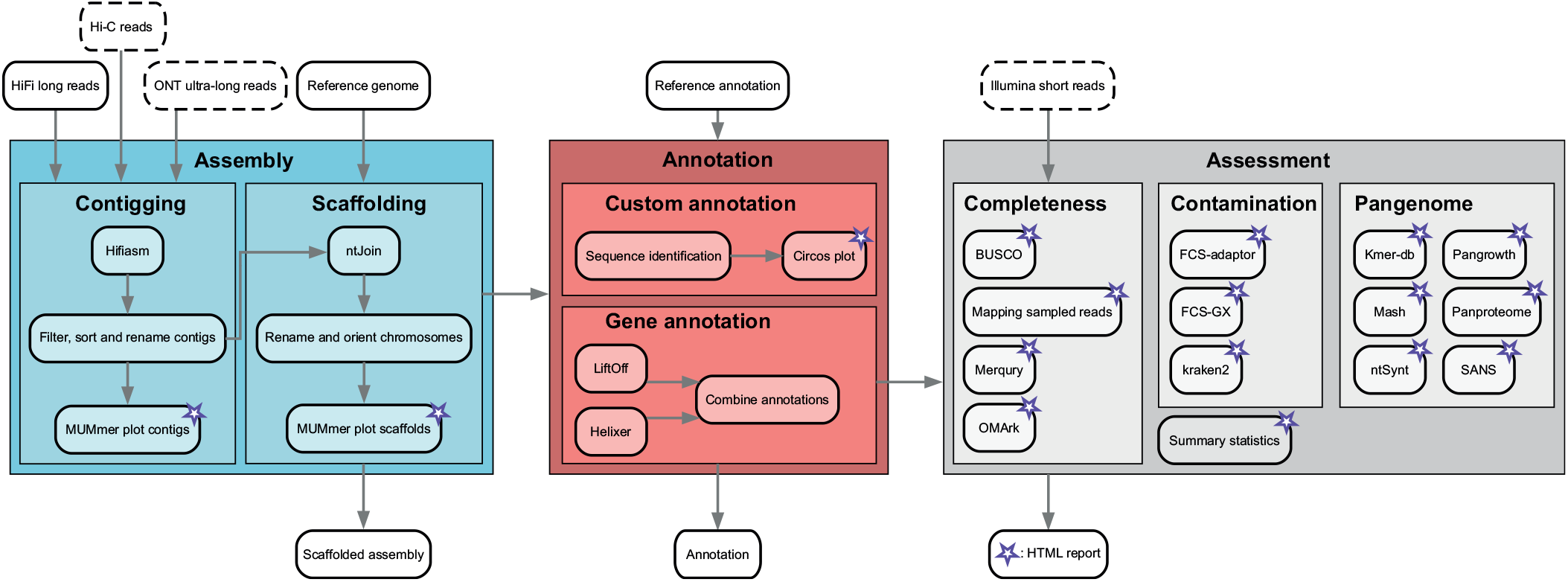
Schematic overview of the MoGAAAP pipeline. The pipeline consists of three main parts, which are subdivided into multiple modules. The main input and output files are highlighted above and below the modules, respectively. Input files in dashed boxes are optional. All rules of the pipeline that create output for the final reporting are highlighted with a purple star.

### Assembly

#### Contigging

Input HiFi reads are assembled using hifiasm v0.19.9 [Cheng et al., 2021] with default parameters and if ultra-long ONT reads are provided, they are passed on to hifiasm using the ‘--ul’ parameter. Hi-C reads, if provided, are also passed to hifiasm. As alternatives to hifiasm, we included both Verkko v2.1 [Rautiainen et al., 2023] and flye v2.9.5 [Kolmogorov et al., 2019], which demonstrates the modularity of the pipeline (future assemblers can be added). Both homozygous and heterozygous diploid genomes are supported by hifiasm, resulting in a phased assembly for heterozygous genomes. Similarly, for Verkko and flye, hapdup v0.12 [Kolmogorov et al., 2023] is used to obtain such a phased assembly. All contigs are retrieved from the resulting assembly graph and written to FASTA format. This FASTA file can be filtered for short contigs and is sorted on size. MUMmerplot v4.0.0rc1 [Marçais et al., 2018] then generates a plot that is added to the report for visual inspection of the resultant assembly compared to a predefined reference genome. The pipeline does not perform filtering of the input sequencing data, so as to reflect the content of the original input library. However, if contamination is detected through the QA module, filtering of input reads may be required to improve the assembly quality. Separation of nuclear and organellar sequences occurs in the “Custom annotation” module.

#### Scaffolding

Because for many species relevant to science and society at least one high-quality chromosome-level genome assembly is now available, we implemented reference-guided scaffolding and chromosomal orientation in the pipeline, useful to efficiently scaffold additional genomes for the same species. For this, MoGAAAP uses the minimizer-based tool ntJoin v1.1.4 [Coombe et al., 2020] with user-provided values for the window and *k* - mer length or alternatively RagTag which is an alignment-based approach to reference-guided scaffolding [Alonge et al., 2022]. The resulting chromosomes are renamed and oriented to be consistent with the supplied reference genome (obtained via a quick mashmap mapping) [Ondov et al., 2016]. MUMmerplot is used again to generate a plot for the report for visual inspection of the scaffolding, renaming and orienting processes. Hi-C reads are not used for automated scaffolding as no currently available tool guarantees correctly scaffolded chromosomes. Instead, if Hi-C reads were provided, a contact map is generated using HapHiC’s plotting function [Zeng et al., 2024] to assess the reference-guided scaffolding.

### Annotation

#### Gene annotation

Accurate and comprehensive *de novo* gene annotation, as would typically be done for a first reference genome for a species, is currently an iterative and time-consuming process, which cannot be fully automated. Therefore, we implemented a workflow that creates a provisional annotation by combining Liftoff v1.6.3 [Shumate and Salzberg, 2021] and Helixer v0.3.2 [Holst et al., 2023] runs on the scaffolded assembly. Since neither tool requires repeat masking, this speeds up the process of gene annotation significantly. For Liftoff, the annotation corresponding to the high-quality chromosome-level reference assembly that was used for reference-guided scaffolding is used. Liftoff tries to transfer all annotated gene features, which may include both protein-coding and non-coding genes. Liftoff is run with the ‘-copies -cds -polish’ parameters to adjust for potentially slightly shifted gene models in the newly assembled genome. Since Liftoff may yield invalid open reading frames, we use AGAT v1.4.0 [Dainat, 2024] to remove these. This Liftoff annotation is then supplemented with Helixer predicted protein-coding genes, again using AGAT; in case of an overlap between Liftoff and Helixer, we favour the Liftoff prediction. Although the quality of this provisional annotation is typically high, it is not evidence-based, necessitating cautious use of the predicted genes. Transcript or protein data can be aligned to estimate the reliability of gene models. Alternatively, if resources are available, an evidence-based annotation can be produced, after which MoGAAAP could be used for quality assessment.

#### Custom annotation

Genome assemblies may be generated for the investigation of specific gene families (e.g. NLRs, transcription factors) or specific sequences (e.g. telomeres, centromeres, rDNA arrays) across genomes. These user-provided queries can be both nucleotide or protein sequences and are searched using BLAST v2.15.0 [Camacho et al., 2009] against the scaffolded assembly. All these queries are used to create a Circos v0.69.9 [Krzywinski et al., 2009] configuration file, which can be polished further to address the user’s needs outside of the pipeline. Additionally, organellar sequences are identified and separated from the ‘nuclear’ assembly based on a hit of *>* 50% of a contig’s length to organellar genomes (separation only for final reporting, not for QA).

### Assessment of assembly and annotation quality

#### Completeness

First, merqury v1.3 [Rhie et al., 2020] is employed for calculating a consensus quality value (QV) based on HiFi reads. Optionally, we allow the user to specify Illumina reads (not used in the assembly process) for an independent check of *k* -mer completeness. MoGAAAP also provides a mapping report of Illumina reads for this. In addition, the pipeline runs two gene completeness assessment tools: BUSCO v5.8.0 [Manni et al., 2021] and OMArk v0.3.0 [Nevers et al., 2024]. BUSCO scores have traditionally been the gold standard for gene completeness; however, in the contemporary era of HiFi assemblies, incomplete assemblies (*<* 90% complete) have become a rarity. Completeness assessed against the larger OMA database is therefore a more informative metric of annotation completeness. Together, BUSCO and OMArk give an insight in gene completeness of the assembly and annotation.

#### Contamination

In addition to OMArk, which summarises possible contamination in the predicted proteome, contamination of the assembly itself is assessed using two separate approaches: kraken2 v2.1.3 [Wood et al., 2019] and FCS-GX v0.5.0 (the NCBI tool for finding potential contamination) [Astashyn et al., 2024]. Kraken2 (ideally run against a recent ‘nt’ database) is used for creating a krona v2.8.1 report [Ondov et al., 2011] showing the likely origin of each scaffold/contig based on *k* -mer composition. Together with the FCS-GX report, this is provided to the user for deciding which scaffolds/contigs need to be removed from the assembly.

#### Pangenome

Finally, we provide insights into a pangenome of user-defined sets of accessions to assess its diversity. Since typically multiple related accessions may be sequenced for assembly, pangenome completeness can provide an overview of the covered diversity of the related accessions. At the genome structure level, we use ntSynt v1.0.0 [Coombe et al., 2024] to calculate collinearity between the assemblies based on minimizers (which makes it independent of gene annotation). The resulting plot can be used to identify large inversions and translocations, indicating potential misassemblies or real structural variation. Next, we employ two separate tools for calculating mash distance between the assemblies: mash [Ondov et al., 2016] and kmer-db v1.11.1 [Deorowicz et al., 2019]. Kmer-db performs the calculation on the entire set of *k*-mers whereas mash subsamples them, affecting runtime and accuracy. The resulting heatmaps give insight into the phylogenetic relationships between the assemblies; this can be compared to the known evolutionary origin of the accessions for identifying potential sample swaps or mislabeling. For assessing the openness of the pangenome, we use both *k* - mer and gene-based metrics. We employ pangrowth (@ commit 71d67bde89326644f6718c82ec2ee7b751f3080b) [Parmigiani et al., 2024] for calculating the pangenome growth based on *k*-mers and PanTools v4.3.1 [Jonkheer et al., 2022] for the pangenome growth based on genes.

## Validation and application

To demonstrate and validate MoGAAAP’s applicability across diverse diploid species, we describe four use cases in Supplementary Information A. First, we showed the usability of the pipeline for the generation of pangenome-sized data by re-assembling and analysing a pangenome of 32 *Arabidopsis thaliana* accessions, for most of which only HiFi data are available [Kang et al., 2023]. Since not all plant genomes are homozygous, we also showed the effectiveness of MoGAAAP on six highly heterozygous grapevine genomes. Next, we demonstrated the applicability of the pipeline to organisms outside of the plant kingdom by assembling a trio from the human pangenome project. Through these use cases, we found that MoGAAAP was able to replicate the findings of the original analyses. Subsequently, we applied the pipeline to a newly generated dataset of *Lactuca serriola*, a wild relative of cultivated lettuce *Lactuca sativa*, on which both HiFi and ONT sequencing have been performed. This resulted in the first chromosome-level genome of *L. serriola*, which we made publicly available.

We found the QA to be a highly important part of the pipeline (see Supplementary Information A). Most assemblies were scaffolded into correct chromosomes and showed little to no contamination. Interestingly, all human genomes were processed correctly, likely because most of the tools employed were developed for the field of human genomics. For the assemblies that showed unexpected outlier statistics in the QA report, we could easily use the report to identify the cause, confirming issues such as contamination and mis-scaffolding based on the output of multiple tools. For example, we found that contamination of foreign DNA was a major factor that influenced the accuracy of the genome assembly and resulting gene annotation. Ideally, all contamination is removed from the input data before assembling the reads. This highlights the iterative nature of the genome assembly process, which we help by providing QA that is as extensive as possible. Therefore, although the pipeline is built to enable an end-to-end automated workflow, some manual curation and filtering remains of vital importance to create a publication-quality genome assembly.

## Strengths and limitations

MoGAAAP is designed to automate and standardise the generation of genomes after the first high-quality genome for a species. It provides a workflow to go from raw input data to a quality assessment report of an assembled and annotated genome, which may be adapted by users to their own needs. Because of the modular structure of the pipeline, it is relatively straightforward to turn a module on or off, or to replace one tool with another. For example, the pipeline employs ntJoin for scaffolding the contigs according to a reference genome. However, for some species there might not be a high-quality chromosome-level reference genome available. In these cases, other methods for scaffolding need to be used instead, e.g. based on Hi-C data. The annotation module where we lift over the genes from a supplied reference genome and add all non-overlapping genes as identified by Helixer provides a similar case: if for the species being assembled, the annotation of the supplied reference genome is of low quality, the resulting annotation will be of low quality too. In this case, it is advisable to run an evidence-based annotation pipeline such as Maker [Cantarel et al., 2008], BRAKER3 [Gabriel et al., 2024] or EGAPx [NCBI, 2024] instead.

The quality of the resulting assembly is dependent on the quality and quantity of the reads supplied. Ideally, the number of contigs should be in the hundreds rather than the thousands. If there are too many small contigs, reference-guided scaffolding using ntJoin will mask structural differences, falsely suggesting full collinearity with the reference genome. Therefore, only structural variation within a single contig compared to the reference genome can be detected. Comparisons of the MUMmerplots before and after reference-guided scaffolding indicates whether this is a potential constraint.

Finally, our pipeline focusses on diploid organisms; however, many organisms have a more complex genetic make-up. Since allopolyploid genomes can be treated as heterozygous diploid genomes, each haplotype is expected to assemble separately. On the other hand, organisms with autopolyploid or aneuploid genomes are currently very challenging to assemble. Since currently no satisfactory assembly tool exists for such genomes, MoGAAAP cannot produce high-quality genomes for them. In the future, it will be relatively easy to include such tools, once available, due to the modular set-up of MoGAAAP.

## Conclusion

In conclusion, we developed a general purpose, scalable and modular pipeline for the assembly of HiFi data, subsequent annotation and quality assessment. In the use cases provided we highlight the importance of QA and subsequent custom curation of the input and/or output, which remains essential to obtain publication-quality genomes. Also, we demonstrated MoGAAAP’s use from single genome assemblies to entire pangenome datasets and from small to large genome sizes.

Pipelines such as MoGAAAP will be increasingly important now that the field of genomics is moving towards pangenomic analyses, which critically depend on the availability of large numbers of related high-quality genomes. A standardised pipeline that can take sequencing data to useful (visual) output in a fast, accurate and comprehensive manner ensures data consistency and quality for downstream applications.

## Supplementary Information A

Use cases for a *L. serriola* genome, an *A. thaliana* pangenome, a grapevine pangenome and a human trio.

## Data availability

The novel *L. serriola* genome, which is described as a use case in Supplementary Information A, has been published at NCBI GenBank (GCA 051521515.1). Underlying HiFi and ONT sequencing data has been published under BioProject PRJNA412928. HTML reports created by MoGAAAP for the use cases described in Supplementary Information A are available through data.4TU (DOI: 10.4121/4b39da65-2eef-4e05-8583-25f8850cf932).

## Competing interests

No competing interest is declared.

## Author contributions statement

D.M.W.: Conceptualization, Data curation, Software, Writing – original draft; K.K.D.: Data curation, Validation, Writing – review & editing; A.K.: Conceptualization, Writing – review & editing; D.L.: Resources; D.R.: Writing – review & editing; M.E.S.: Writing – review & editing; R.W.M.: Conceptualization, Supervision, Writing – review & editing; S.S.: Supervision, Writing – review & editing.

## Acknowledgments

The authors thank Huaqin Xu for helping with the genome submission to Genbank and Keri Cavanaugh for preparing the HiFi libraries for *L. serriola*.

## Funding

This work was supported in part by the LettuceKnow project (with project number 1.1B) of the research Perspective Program P19-17 which is (partly) financed by the Dutch Research Council (NWO) and the breeding companies BASF, Bejo Zaden B.V., Limagrain, Enza Zaden Research & Development B.V., Rijk Zwaan Breeding B.V., Syngenta Seeds B.V., and Takii and Company Ltd and the International Lettuce Genomics Consortium 3, which is funded by the Crops of the Future Program of the Foundation for Food and Agricultural Research with matching funding from 14s breeding companies.

## Notes

https://doi.org/10.4121/4b39da65-2eef-4e05-8583-25f8850cf932

### Competing Interest Statement

The authors have declared no competing interest.

