## Supplementary Information for "MoGAAAP: A modular Snakemake workflow for automated genome assembly and annotation with quality assessment"

### Motivation

Across plant, animal and human genomics, population studies increasingly focus on obtaining high-quality genome assemblies for pangenomic studies of diversity panels. Typically, these species already have one or more reference genomes available, simplifying the generation of successive genomes for the same species based on long-read sequencing techniques such as PacBio HiFi sequencing. MoGAAAP was designed to precisely address this challenge: automate the generation of an annotated chromosome-level genome assembly with an extensive QA report for species with already one high-quality genome available.

The development of MoGAAAP was sparked initially by a sequencing project in lettuce, aiming to deliver high-quality assemblies of many genotypes, complementing a handful of available high-quality references. As similar efforts are made for many important plant species, animal species and human populations, there is great potential for a standardised workflow. While we designed the basis of MoGAAAP to assemble and annotate lettuce genomes in high throughput, we realised that the general design makes it applicable to all such eukaryotic species.

Here, we present four different use cases to demonstrate the functionality of the MoGAAAP pipeline (github.com/dirkjanvw/MoGAAAP). First, we validate the working of MoGAAAP on three use cases (two plant datasets and one human), after which we apply it to one of our lettuce genomes: *Lactuca serriola*. For reproducibility, we provide the input and output (HTML report) for each use case in a data repository at data.4TU (DOI: 10.4121/4b39da65-2eef-4e05-8583-25f8850cf932).

### Validation

#### *Arabidopsis thaliana*

To showcase the ease of using MoGAAAP for pangenome studies, we reproduced a pangenome set of assemblies using our pipeline. Pipelines for the automated assembly, annotation and assessment are especially useful for large-scale sequencing projects to reduce the number of technical artefacts between the resulting assemblies. Here, we re-analysed the 32 *Arabidopsis thaliana* pangenome from Kang et al. (2023).

The publicly available HiFi data for all accessions were downloaded for assembly, as well as the Illumina data for QA of two accessions (CNCB BioProject PRJCA012695) and Hi-C data for one accession (SRR20242239). All accessions were assembled into 51 to 819 contigs and the pipeline was able to scaffold and rename all but one accession relative to the five chromosomes of the *A. thaliana* TAIR10 assembly. The resulting assemblies had a QV between 50 - 70 given their input HiFi data, indicating high accuracy. Annotation by LiftOff and Helixer resulted in 32,711 - 47,793 predicted gene models with a BUSCO completeness between 91.9% and 99.9%.

Although the resulting assemblies were complete and largely congruent with expectations, they were not identical to the findings of Kang et al. (2023). Three accessions (Yilong, SB30 and Tibet) had notably larger numbers of contigs and predicted genes, which seemed to correlate with the amount of potential contamination in these assemblies (as revealed by the Kraken2 and FCS-GX analyses) (Supplementary Figure 2). Only unscaffolded contigs were labelled as contaminated, and the total number of genes on only the five chromosomes per accession was 29,663 - 34,621. This number closely matched the expected number of genes in ARAPORT11 (33,243), indicating that the scaffolds themselves were likely free of contamination. Therefore, it is likely that Kang et al. (2023) performed contamination filtering prior to assembly, though undocumented.

The assemblies from MoGAAAP could not immediately be used for a pangenome study, because they still require human correction. For example, analysing the MUMmerplots to TAIR10 in the report (Supplementary Figure 1), it was evident that there was a misassembly of chromosome 2 in Col-0 that originated in the hifiasm assembly and had been propagated through the scaffolding phase. Comparison of the MUMmerplot and Hi-C plot for Col-0 confirmed the incorrect assembly for this contig. Such inconsistencies need to be manually resolved by 1) cutting the assembly and manually joining the correct sequences together, or 2) changing the assembly algorithm. Changing the assembly algorithm to another assembler or by trying a range of parameter changes takes more time, making the manual correction based on e.g. Hi-C evidence more attractive. In summary, the automated QA statistics and figures were very helpful in finding and solving mistakes in an informed manner.

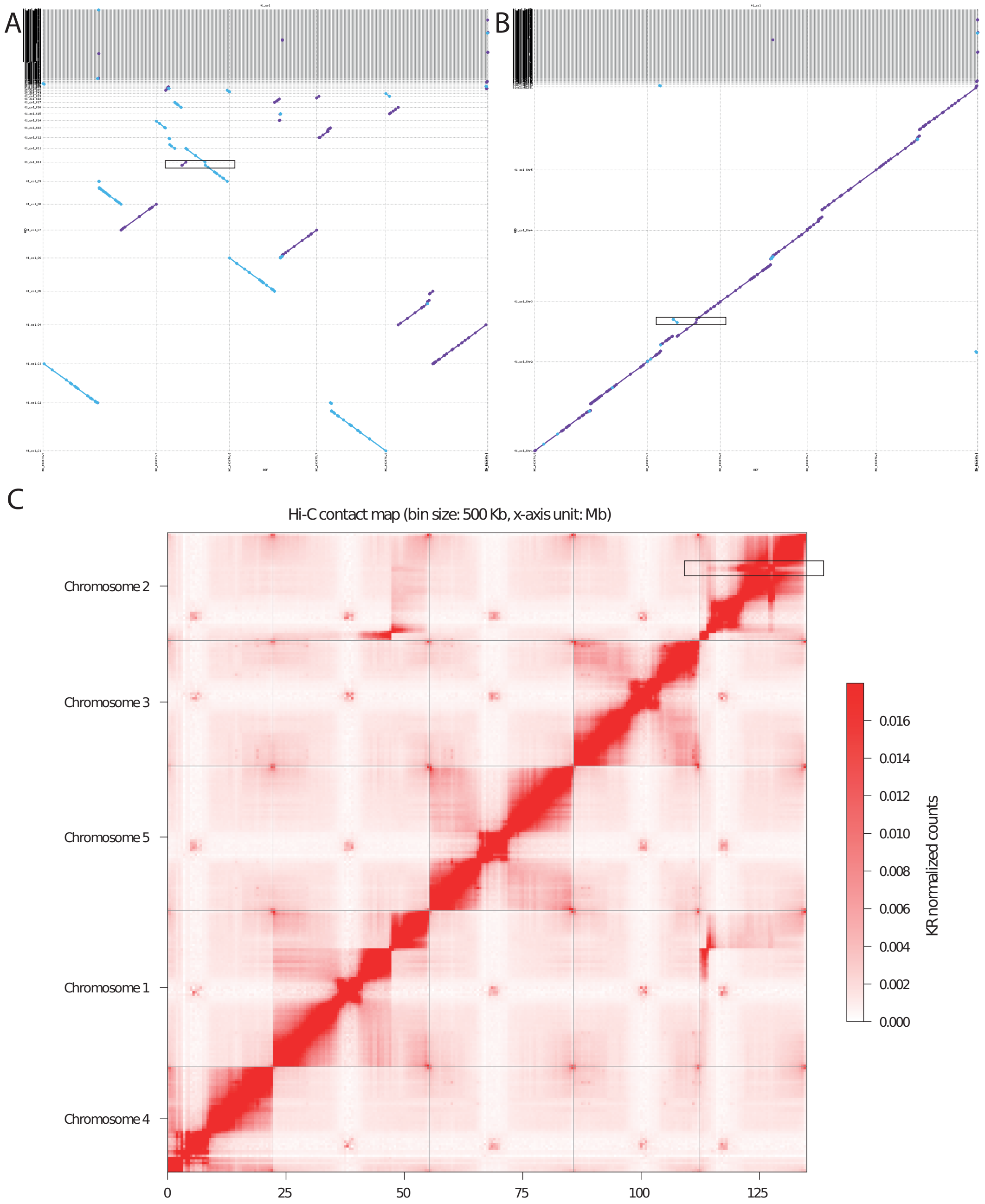

Supplementary Figure 1: Evidence of a misassembly by hifiasm in the *A. thaliana* Col-0 contigs for chromosome 2 propagated to the scaffolded assembly. The misassembly in part of chromosome 2 is highlighted in the figure by the black rectangle. Three plots provide evidence for this: A) MUMmerplot of the hifiasm contig assembly of Col-0 against the public Col-0 TAIR10 reference genome. B) MUMmerplot of the scaffold assembly against the public Col-0 TAIR10 assembly. C) Hi-C heatmap for the scaffolded assembly: numbers on the x-axis indicate the cumulative length of the chromosomes in mega basepairs.

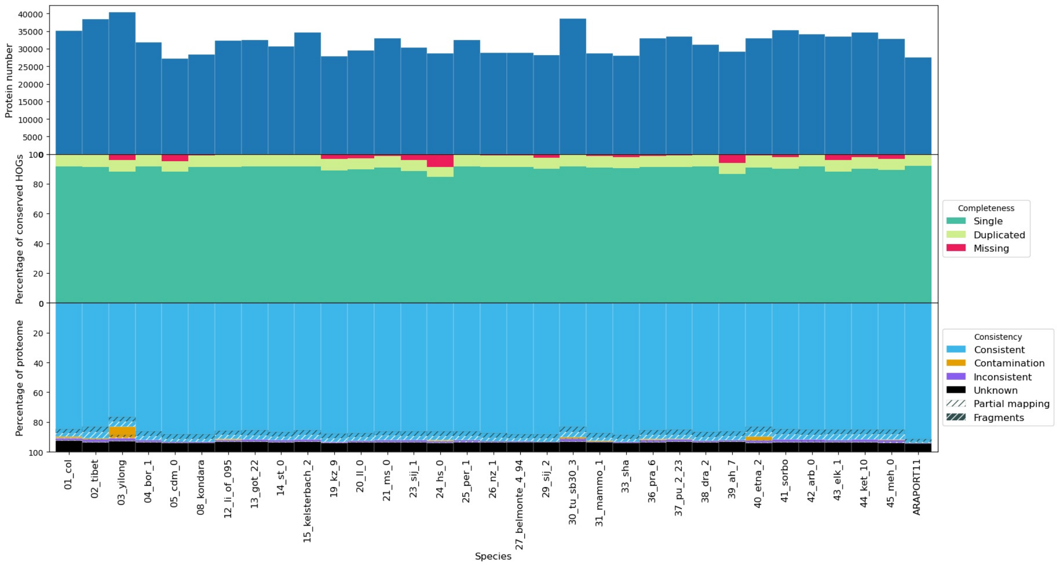

Supplementary Figure 2: OMArk result of analysing the proteomes of the provisional annotations for the 32 *A. thaliana* created by MoGAAAP. Most proteomes are very complete and consistent, and have about the same number of ∼30,000 proteins. However, some proteomes that were larger showed clear indications of contamination (therefore, genes predicted in non-*A. thaliana* sequences).

#### Grapevine

Grapevine is a highly heterozygous diploid plant species that was recently used for a pangenome study in which novel accessions were sequenced with HiFi, ONT and Hi-C (Liu et al., 2024). We assembled six publicly available accessions using MoGAAAP employing all three data types, and succeeded in obtaining the separate haplotypes from the hifiasm assembly.

All chromosome statistics closely matched the values reported by Liu et al. (2024) without any manual correction. The chromosomes were fully phased by the hifiasm assembler and had a QV of 50 to 65, indicating high accuracy. However, the comparison also revealed some differences. The MUMmerplots identified a potential scaffolding error where two chromosomes had been joined in one of the two Hongmunage haplotypes, which was confirmed in the Hi-C contact map. This difference is likely caused by a different scaffolding algorithm: Liu et al. (2024) used RagTag whereas MoGAAAP uses ntJoin by default. Also, the BUSCO score for one of the Shine Muscat haplotypes from MoGAAAP was found to be 0. Looking into this problem, it turned out to be caused by the BUSCO software as Helixer had annotated a protein larger than 100,000 amino acids, which is not allowed by BUSCO.

Upon closer inspection of the QA report from MoGAAAP, some contamination problems of the grapevine reads became obvious that were previously not reported by Liu et al. (2024). Hongmunage was also the only accession with a large amount of contamination as identified by FCS-GX, Kraken2 and OMArk. This was also reflected by the large number of genes on haplotype 1 (49,375) of this accession versus all other grapevine haplotypes (∼31,000 to ∼39,000) as well as the large amount of unscaffolded sequence. According to FCS-GX and Kraken2, the main contaminants were *Erisyphe necator* (grape powdery mildew) and insect. Interestingly, the contamination was not mentioned by Liu et al. (2024) even though a clean assembly was reported without a significantly larger number of genes or unscaffolded contigs. The amount of contamination by grape powdery mildew was so large that it showed up in the k-mer spectra plots as well, which indicated it was assembled in almost its entirety as part of haplotype 1 of Hongmunage (Supplementary Figure 3). The best way to resolve this issue would therefore be to clean the reads of Hongmunage of *Erisyphe necator* and insect reads and assemble it again, but to keep the assemblies and annotations as generated by MoGAAAP for all other accessions. Cleaning the reads of Hongmunage could for example be done by aligning them to the contaminant assemblies or running kraken2 on the reads and filtering out all contaminant reads.

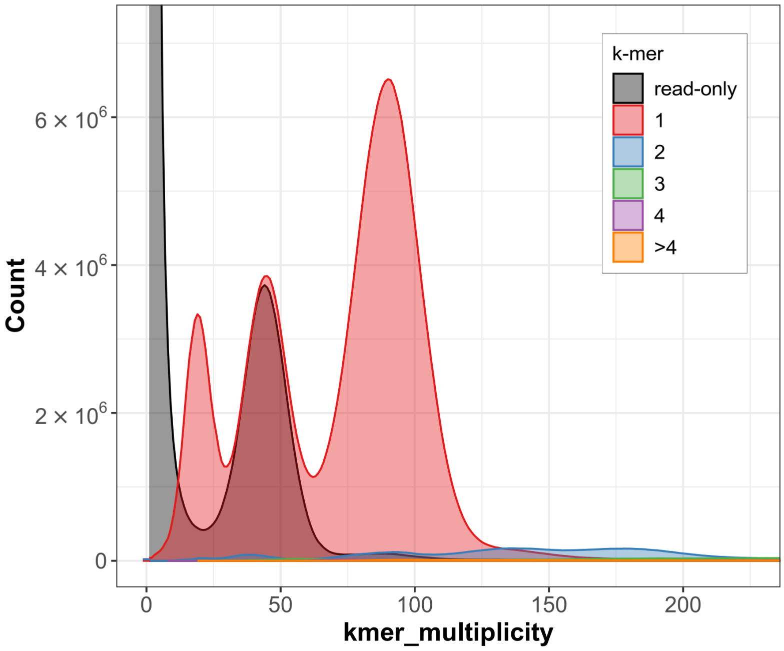

Supplementary Figure 3: K-mer spectra-CN plot created by merqury for the HiFi reads of grapevine cultivar Hongmunage against the haplotype 1 assembly of this accession. Strangely, three red peaks (showing HiFi k-mers that occur once in the assembly) are visible instead of the expected two. The right-most peak represents the k-mers present in both haplotypes of Hongmunage, the middle peak represents the k-mers that are haplotype-specific in Hongmunage (at half multiplicity compared to the right-most peak), and the left-most peak represents the k-mers of a large contaminant species present in the HiFi data (likely grape powdery mildew).

#### Human trio

MoGAAAP is not only designed for plants but for any eukaryotic genome. In order to demonstrate applicability to non-plant species, we applied the pipeline to a human dataset. With the rise of pangenomics, it is important to have a pipeline that can ensure consistency in assembly and structural annotation. We therefore assembled the HiFi data of one of the trios of the human pangenome project: HG002, HG003 and HG004. In addition, for HG002, we also used Hi-C data to show its integration in the pipeline.

After hifiasm assembly, reference-guided scaffolding with ntJoin was able to put 95.5% - 97.5% of the assembled length into chromosomes based on the GRCh38 reference genome for each haplotype. QA was performed on all haplotypes separately. Confirming the accuracy of the scaffolding, the MUMmerplot figures showed full collinearity for all chromosomes; except for the mother HG004 which lacked a Y chromosome as expected, and X chromosomes of the son and father which were not always assembled correctly (also compare to the contig-level MUMmerplot). Furthermore, the contact map generated with the Hi-C data showed no miss-scaffolding for HG002.

The QV as calculated by merqury based on the HiFi data showed high k-mer completeness (66.0 - 67.3) for each haplotype. Correspondingly, BUSCO and OMArk showed full completeness of the assembly and annotation (95% - 98% and 94% - 97%, respectively). Confirming previous work on these cell lines, only human gammaherpesvirus 4 (also known as Epstein-Barr virus (EBV)) contamination was found in the assemblies using both Kraken2 and FCS-GX. EBV was used to immortalise the cells and therefore expected to still be present genomically (Zook et al., 2016). This analysis shows that non-plant species may also be assembled, scaffolded, analysed, annotated and assessed for quality using MoGAAAP.

### Application

To show that MoGAAAP, in combination with manual curation, can be used for the creation and assessment of a publication-quality genome, we applied it to a novel *Lactuca serriola* genome.

#### *Lactuca serriola*

*L. serriola* is the likely progenitor of cultivated lettuce (*Lactuca sativa*) and is of major importance for lettuce breeding as a source of wild alleles (Wei et al., 2021). No high-quality genome assembly has been published for *L. serriola*. Therefore, we used our pipeline to assemble the genome of *L. serriola* (US96UC23) using HiFi and ONT reads and scaffold against a high-quality *L. sativa* genome as the two species are closely related.

##### Sequencing information

*L. serriola* (US96UC23) seedlings were grown in Magenta boxes under sterile conditions in the dark. Seven day old etiolated seedlings were harvested and flash frozen in liquid nitrogen. Etiolated tissue was used for DNA extraction using a CTAB DNA extraction/sorbitol cleanup protocol. The HiFi library was prepared using PacBio SMRTbell prep kit 3.0 followed by bead clean up and size selection (>13.6 kb). HiFi sequencing was performed using the PacBio Revio platform aiming for 20x coverage. In total 56.9 Gb of HiFi reads were generated. ONT libraries were prepared with the LSK109 kit. ONT sequencing was performed using PromethION on an R9.4.1 chip aiming for 140x coverage and base called with Guppy (v5.0.16.sup). 10.8M reads (total 248.8 Gb) were generated, after which Porechop v0.2.3 was used to remove residual ONT adapters and NanoFilt v2.7.1 was used to select reads with an average quality score >Q10.

##### Genome assembly

Genome assembly was performed using hifiasm with all HiFi reads and a subset of the ONT reads, specifically only reads > 36kb. In total, we used 56.9 Gb of HiFi reads and 84.0 Gb of ultra-long ONT reads, corresponding to an expected coverage of 21.9x and 32.3x coverage respectively. The resulting contig assembly size of 2.559 Gb was consistent with the expected genome size of 2.6 Gb. In order to scaffold the 258 contigs relative to the *L. sativa* Salinas v11 RefSeq genome (GCF_002870075.4) (Reyes-Chin-Wo et al., 2017), we had to relax the default ntJoin parameters for window length to 40,000 bp (with $k = 52$), because *L. serriola* is a different species than *L. sativa*. Failing to do so caused chromosome joining. Scaffolding resulted in 29 contigs scaffolded into nine pseudomolecules, making up 99.35% of total assembly length. The remaining 229 contigs (0.65% of total assembly length) could not be scaffolded. Of these, 92 were identified as organellar contigs.

##### Annotation

Next, the pipeline performed the lifting over of genes from v11 of *L. sativa* Salinas RefSeq genome annotation. LiftOff was able to transfer the coordinates of 40,254 genes (of which 29,693 were protein-coding). This was supplemented with 9,283 non-overlapping gene models from Helixer. In total, this resulted in 38,976 protein-coding genes. This is in the same range as other annotated *Lactuca* sp.: *L. sativa* Salinas has 36,855, *Lactuca saligna* CGN05327 has 42,908 and *Lactuca virosa* CGN04683 has 39,887 protein-coding genes annotated.

##### Quality assessment

Based on an independent Illumina short-read sequencing dataset (CNS0047707) of the same accession by Wei et al. (2021), the QV value of the assembly was 50.5 (separate chromosomes ranging from [52 - 56]) and the assembly was 99.06% complete. Also, the spectra-CN plot with the independent Illumina short-read data confirmed a correct assembly: no read- or assembly-only k-mers were identified. BUSCO and OMArk scores confirmed the high completeness of the assembly and annotation with values of 98.2% and 97.8% respectively. The extensive contamination screening by the pipeline (OMArk, Kraken2, FCS-GX) was not able to find any non-lettuce sequence, highlighting the robust library preparation. Interestingly, a large inversion on the right arm of chromosome 1 with respect to *L. sativa* Salinas v11 was identified in the MUMmerplot. The location of this ∼24Mb inversion was consistent with the genetic map previously published by Reyes-Chin-Wo et al. (2017) (see their Supplementary Figure 9) which is a cross between *L. sativa* Salinas and the here assembled *L. serriola* US96UC23. For validation, we used the alignment of both HiFi and ONT long-read data to confirm the correctness of the assembly (Supplementary Figure 4).

##### GenBank submission

The scaffolded assembly and annotation were prepared for submission to GenBank, which involved file formatting and a final confirming that they were of high quality (free of contamination, complete and accurate). During this iterative process, both assembly and annotation had to be slightly adjusted. We used GAG (Geib et al., 2018) to convert the files to tbl format and identify potential issues. For the assembly, we took the nuclear assembly and shortened the sequence headers. For the annotation, we first renamed all genes to remove traces of the gene identifiers from *L. sativa* Salinas v11, retained only the longest isoform per gene and removed all non-coding genes. Some genes had overlapping CDS sequences on opposing strands; for these, the shortest of each pair was removed. Finally, we removed all genes without start or stop codons and all genes with introns smaller than 10 nucleotides because these are not permitted by GenBank. After the cleaning process, 37,518 protein-coding genes remained.

The *L. serriola* assembly can be found at GenBank GCA_051521515.1. The underlying HiFi and ONT reads have been released under BioProject PRJNA412928.

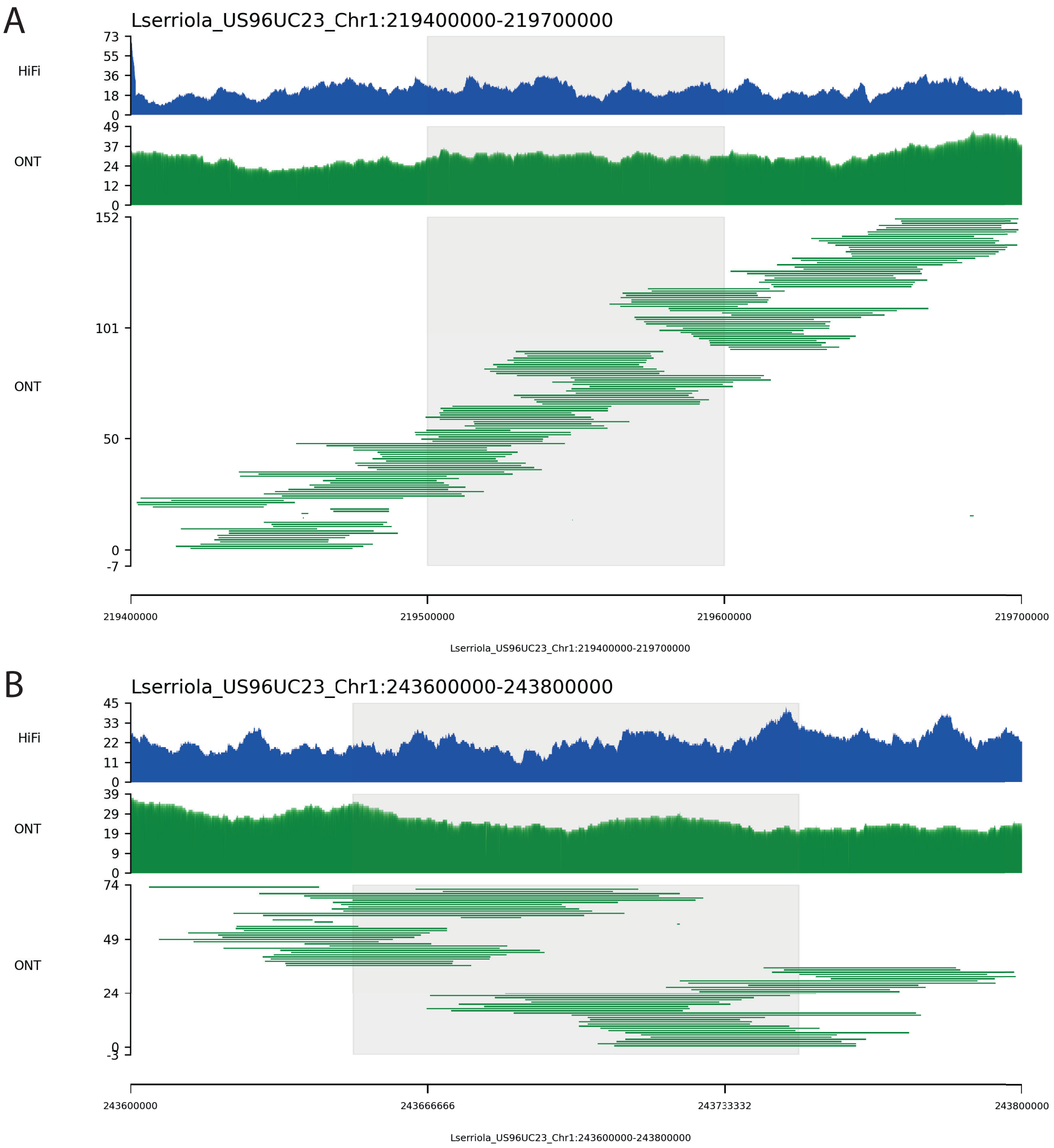

Supplementary Figure 4: The left (A) and right (B) side of the ∼24Mb inversion of *L. serriola* UC96UC23 compared to *L. sativa* Salinas v11 (the location of the breakpoint is highlighted in gray). Coverage is plotted for both HiFi and ONT (top two tracks) and all individual ONT reads fully within the region are shown in the third track. Since neither HiFi nor ONT shows any drop in coverage or breakpoint, the *L. serriola* assembly is structurally correct and the inversion real.
